# Mating disrupts morning anticipation in *Drosophila melanogaster* females

**DOI:** 10.1101/2022.05.18.492405

**Authors:** Sabrina Riva, Juan Ignacio Ispizua, María Trinidad Breide, Sofía Polcowñuk, José Ricardo Lobera, M. Fernanda Ceriani, Sebastian Risau Gusman, D. Lorena Franco

**Affiliations:** Medical Physics Department, Bariloche Atomic Centre Comisión Nacional de Energía Atómica (CNEA) and Consejo Nacional de Investigaciones Científicas y Técnicas (CONICET), San Carlos de Bariloche, Argentina; Laboratorio de Genética del Comportamiento. Fundación Instituto Leloir - IIBBA - CONICET, Buenos Aires, C1405BWE, Argentina; Institute of Cancer Sciences, University of Glasgow, G61 1QH Bearsden, Scotland

**Author notes:** These authors contributed equally to this work (Sabrina Riva, Juan Ignacio Ispizua). Shared corresponding authors.

## Abstract

After mating, the physiology of *Drosophila* females undergoes several important changes, some of which are reflected in their rest-activity cycles. To explore the hypothesis that mating modifies the temporal organization of locomotor activity patterns, we recorded the fly activity by a video tracking method. Monitoring rest-activity patterns under light/dark (LD) cycles indicated that mated females lose their ability to anticipate the night-day transition, in stark contrast to males and virgins; this postmating response is mediated by the sex peptide (SP) acting mainly on *pickpocket* (*ppk*) expressing neurons, since reducing expression of the SP receptor (SPR) in these neurons restores the ability to anticipate the LD transition in mated females. We further provide evidence of connectivity between PPK+ neurons and the pigment-dispersing factor (PDF)-positive ventral lateral neurons (sLNv), which play a central role in the temporal organization of daily activity. Since PDF has been associated to the generation of the morning activity peak, we hypothesized that the mating signal could modulate PDF levels. Indeed, mated females have reduced PDF levels at the dorsal protocerebrum; moreover, SPR downregulation in PPK+ neurons mimics PDF levels observed in males. In sum, our results are consistent with a model whereby mating-triggered signals reaches clock neurons in the fly central nervous system to modulate the temporal organization of circadian behavior according to the needs of the new status.

**Author Summary:** After mating, *Drosophila* females undergoes striking behavioral changes, specially in their activity patterns. Despite some of the circuits that deliver mating signals to the female brain are known the connection with the circadian network has not been explored in detail. Here, we show that mating changes the onset of daily activity, masking a central function of the clock. This modulation is mediated by the sex peptide (transferred during courtship) acting on PPK+ neurons, which, in turn, directly contact PDF+ neurons, responsible for the increase of the activity that precedes dawn. Thus, our work identifies a postmating response directly related to the circadian clock, and begins to unravel the underlying neuronal circuit.

## Introduction

In most animals, endogenous circadian clocks coordinate physiological and behavioral processes to keep the entire organism in synchrony with the 24 hours day-night cycling environment. In *Drosophila melanogaster* the circadian clock is composed by approximately 150 neurons, where the steady-state levels of clock-related proteins oscillate with periods close to 24 hours. These neurons are organized in different clusters (1,2) whose interaction is necessary for a coherent and plastic control of behavior (3). The ventrolateral neurons (LNvs) are important because they drive rhythmicity under free-running conditions (4–7), mostly through the release of *pigment dispersing factor* (PDF), a neuropeptide expressed in the small (sLNvs) and large LNvs (lLNvs) which is key for the communication between them and other group of the clock neurons (5,8).

In many species, mating induces changes in the behavior and physiology of females; these changes are known as postmating responses (PMR). In *Drosophila* females, mating induces a reduction in partner receptivity (9,10), an increase in egg production and oviposition (11), stimulation of the immune response (12) and changes in sleep and activity patterns (13,14), as well as in nutritional preferences (15). These PMRs are mediated by the sex peptide (SP), transferred from males to females during mating (16,17). SP acts mostly via the sex peptide receptor (SPR), on a small number of uterine Sex Peptide Sensory Neurons (SPSN) that co-express the sex-determination genes *doublesex* (*dsx*), *fruitless* (*fru*), and *pickpocket* (*ppk*), triggering various PMRs (17–19). It has also been suggested that SP could act via the haemolymph (20), which is consistent with SPR being expressed broadly in the CNS (21).

Fruit flies display two activity bouts at dawn (the so-called “morning peak”) and dusk (“evening peak”). The early start of the morning activity peak shows that flies can anticipate the transition between night and day. Mated females usually have greater activity than virgin females and males, and less daytime sleep (13,14), a change modulated mainly by SP through SPR within SPSN and Sex Peptide Abdominal Ganglion (SAG) neurons (22). This PMR has also been observed in females of other *Drosophila* species such as *D. suzukii* (23). In general, sleep is not only regulated by the circadian clock, but also by homeostasis (through a 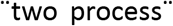 model, Borbély 1982), even in *Drosophila* (24). Thus, the reduction of daytime sleep displayed by mated females could be due to an interference with any of these two processes. An important question is whether the mating status can influence the circadian clock to modify the temporal behavior of mated females.

We show here that mated females lose their ability to anticipate the night-day transition, a clear output of the circadian clock. This PMR is mediated by SP acting on PPK+ neurons, since decreased expression of SPR in these neurons restores morning anticipation in mated females. We searched for projections of *ppk*+ expressing neurons near circadian clusters, and found that the PDF+ sLNvs are postsynaptic targets of PPK+ SPR+ neurons. This connection, along with the relationship between PDF and morning anticipation (4), suggested that PPK+ neurons are involved in the inhibition of either expression or transport of PDF. Accordingly, PDF levels are reduced in the dorsal termini of mated females, which can be partially rescued through SPR expression in PPK+ neurons. Thus, our results are consistent with a model whereby mating-triggered signals are delivered to the clock network in order to modulate the temporal organization of the behavior.

## Results

### Mating induces loss of morning anticipation in mated females

Locomotor activity has been extensively studied in males, but much less in female (mated or virgin) flies, and most published studies were performed using the Drosophila Activity Monitor tracking system (DAM, Trikinetics). On this system, flies are housed in small tubes (¬320 mm^3^), and their movement is recorded only when they cross an infrared beam. It has been reported (25) though, that activity records differ between DAM and the more accurate video tracking systems. Moreover, activity of mated flies depends on the size of the arena where they are placed (25). Thus, in order to provide mated females with a more adequate environment where movement can be accurately recorded, we developed a system where flies are placed in a set of relatively large transparent chambers (80 x 8 x 8 mm), and are tracked using a video system (for a detailed description please refer to the Methods section). In order to validate our setup, we compared the activity recorded by our video tracking method, which directly measures activity as distance travelled per second, with recordings obtained using the DAM system, which estimates activity as the number of interruptions of a single infrared beam, firstly in male flies. Both methods show that flies display bimodal locomotor activity patterns over several 12 h: 12 h LD cycles, and no significant differences were found in the morning anticipation activity between both systems (**Figure S1 A, B, C**). However, a close examination of their sleep pattern revealed that flies sleep significantly more when locomotor activity is monitored using the DAM system than when they are registered using our video tracking method (**Figure S1D, E**). This is probably due to the fact that the DAM system fails to detect a portion of the activity that flies display along the day, for instance when they stand on one side of the tube, without crossing the infrared beam, likely resulting in an overestimation of sleep duration. These results confirm that even though single-fly recordings generated either by the DAM system and our system produce highly similar locomotor activity profiles, validating our experimental setup, video tracking acquisition provides more accurate register of the activity patterns.

Next, we compared the locomotor activity of males, virgin and mated females of different control laboratory lines, *white (w^1118^), Oregon R* (*OreR*) and *Canton-S* (*CS*) under 12 h: 12 h LD cycles. **Figure 1A** shows the average activity profile (AAP) for males, virgin and mated females of *w^1118^* flies. Male flies have two pronounced peaks of daily activity that start in anticipation to the lights-on and lights-off transitions, separated by a relatively long interval with very low activity (siesta). In contrast, mated females displayed a sustained and robust activity during daytime while at night the activity remains as low as in males. Virgin females, on the other hand, are more active than males during daytime but less so than mated females (13,14). A closer visual inspection of AAPs showed that males and virgin females have a gradual increase in activity before lights-on (termed morning anticipation), whereas mated females did not display this morning anticipation, but they had a high amplitude morning startle response coincident with lights-on (**Figure 1B**). A more quantitative assessment of the degree of anticipation is given by the morning anticipation index (MAI, defined in Materials and Methods) (26). **Figure 1C** shows a significant reduction in MAI in mated females compared with males supporting a strongly dimorphic trait. Additionally, the figure shows that mated females display a significant reduction in MAI when compared to virgin females, to the point that morning anticipation is virtually suppressed (on average) suggesting a postmating effect. Interestingly, all three control strains showed a reduced morning anticipation in mated females, supporting the idea that this effect occurs across different genetic backgrounds. Thus, our data shows that one of the postmating responses characteristic of mated females is the suppression of their ability to anticipate the night-day transition. In this article we focused on characterizing this postmating behavior.

**Figure 1.**
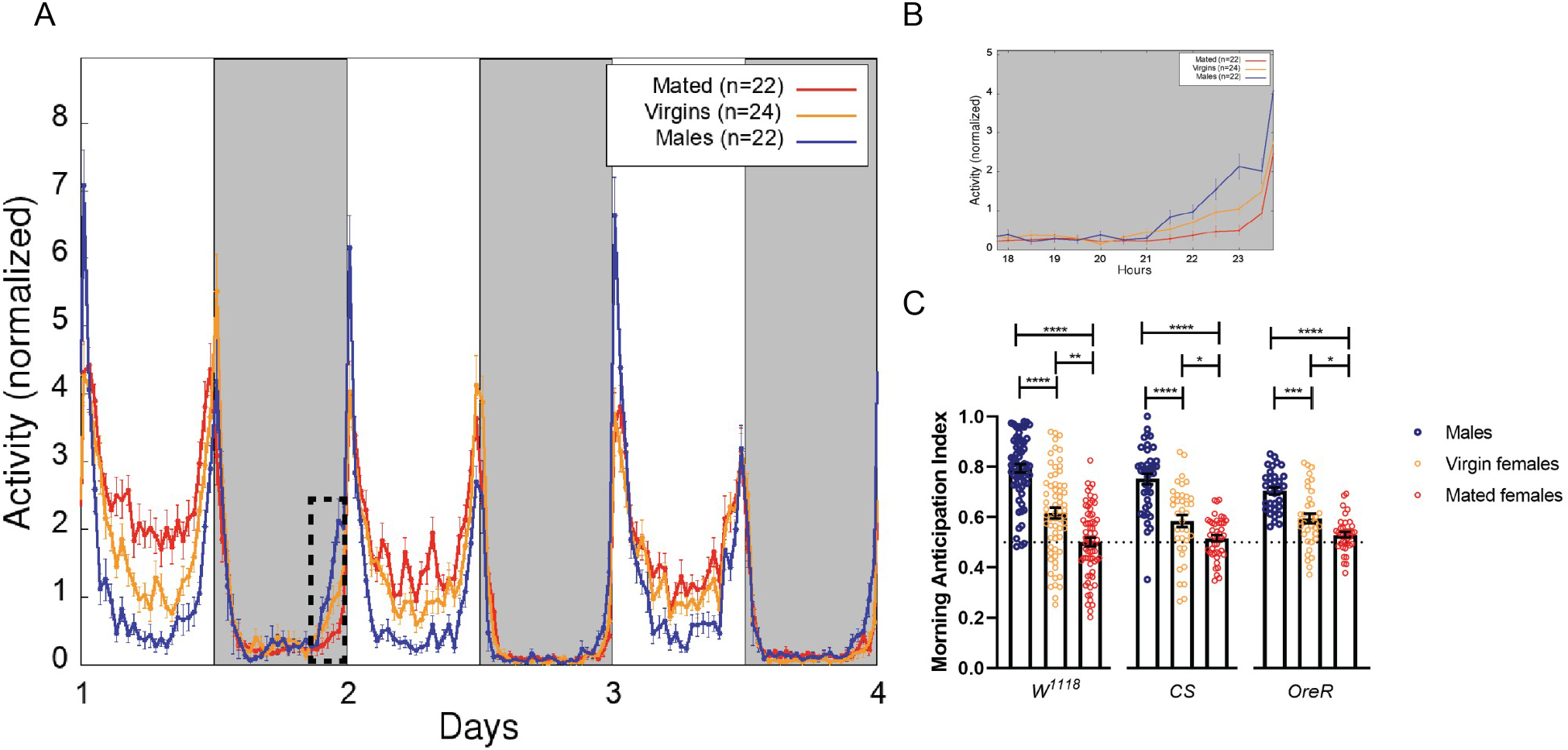
Mating suppresses morning anticipation in females of *Drosophila melanogaster*. ***A***, Average locomotor activity profile of *white (w^1118^)* flies in LD. ***B***, Zoom of the region enclosed by dashed lines in panel A to highlight the difference between groups. ***C***, Anticipation indexes for males (blue), virgin (orange) and mated (red) females of *white, CantonS and Oregon* flies. For *w^1118^* flies: males (n=62), virgin females (n=64) and mated females (n=66). For CS flies: males (n=39), virgin females (n=37) and mated females (n=43) and for Oregon flies: males (n=34), virgin females (n=35) and mated females (n=37). Each dot corresponds to the average index of the three complete days calculated for a single fly. The mean and SEM are shown as bars. Statistical analysis: Kruskal–Wallis test; for the MAI of white flies, X^2^ = 78.02, p< 0.0001; for the MAI of *CS* flies, X^2^ = 49.48, p< 0.0001; for the MAI of *OreR* flies, X^2^ = 44.43, p< 0.0001. Pairwise comparisons were performed using Dunn’s multiple comparisons test. Statistically significant differences are represented by *p<0.05, **p<0.001, ***p<0.0002, ****p<0.0001.

### Loss of morning anticipation is mediated by sex peptide in PPK+ neurons

Although SPR is detected broadly on the female reproductive tract, the ventral nerve cord and the brain (21), only a restricted subset of *fru+/ppk+/dsx+* sex peptide sensory neurons (SPSN) expressing SPR in the reproductive system are necessary and sufficient for inducing SP-mediated postmating responses (17–19). To ascertain whether SPSN neurons were involved in the modulation of morning anticipation through SP signaling, RNA interference (RNAi)-mediated SPR knock down (21) in this group of neurons was used, in order to evaluate its effect on locomotor activity in mated females. Surprisingly, SPR downregulation in SPSN neurons induced a significant increase in morning anticipation compared with mated controls (**Figure 2A**). We next explored the individual contribution of *fru+, ppk+* and *dsx+* neurons in the modulation of morning anticipation. SPR knock down in *fru+* and dsx+ neurons resulted in a significant increase in the MAI compared to mated controls (**Figure 2B and 2C**). This increase in MAI index was similar to the value obtained when SPR expression was reduced in SPSN neurons. However, SPR knock down in PPK+ neurons resulted in a dramatic increase in this index (**Figure 2D**) suggesting a distinctive role of PPK+ neurons in the control of this postmating response. Remarkably, we did not observe any significant difference in morning anticipation when we downregulated SPR in PPK+ neurons in virgins females. Since all virgin genotypes have morning anticipation, **(Figure 2E)** the regulation by SPR is indeed mating-state-dependent. These findings demonstrate the need for SPR function in SPSN neurons to respond to and transmit SP-generated signals to control morning anticipation in mated females, further confirming the importance of these neurons for a variety of post-mating responses (17–19). Data presented herein shows that additional SPR+ PPK+ neurons beyond the SPSN domain play a role in the modulation of morning anticipation. Since connectivity, from the SPSNs to the central brain was previously characterized (17–19,27), we wondered how these additional PPK+ neurons would transmit postmating signals involved in morning anticipation.

**Figure 2.**
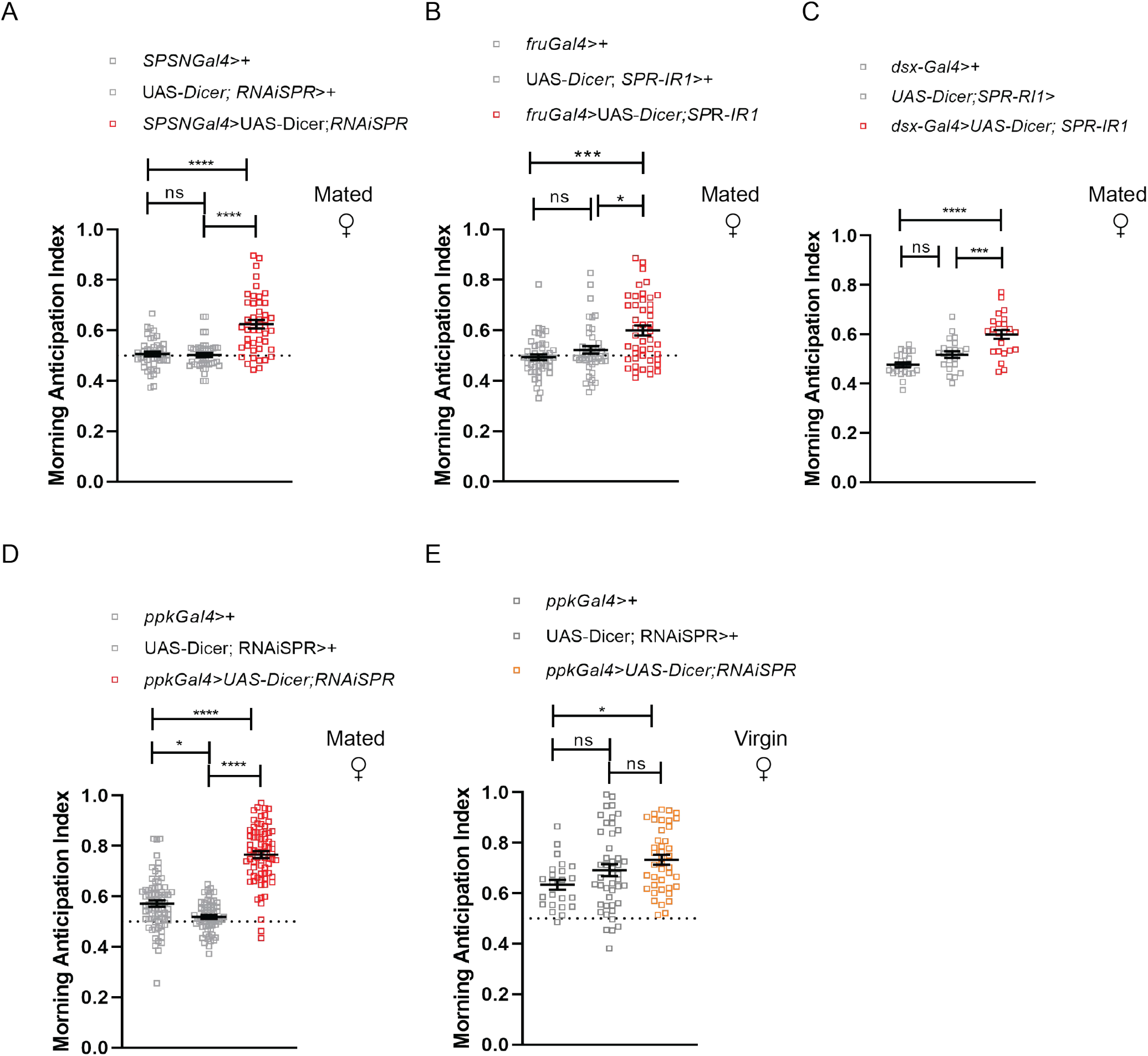
Downregulation of SPR in PPK+ neurons restores morning anticipation. Morning anticipation index in mated females upon chronic SPR knock down in different groups of neurons and their controls. ***A*** *SPSN-Gal4>UAS-Dicer2, SPR-IR*1 (n=45); *SPSN-Gal4*>+ (n=44); *UAS-Dicer2, SPR-IR1*>+ (n=44). Statistical analysis: Kruskal-Wallis test X^2^=36.50 *p* < 0.0001. Pairwise comparisons were performed using Dunn’s multiple comparisons test. Statistically significant differences are represented by ****p<0.0001. ***B*,** *fru*-Gal4>UAS-*Dicer*2, *SPR-IR*1: (n=46), *fru-*Gal4>+ (n=46); UAS-*Dicer2*, *SPR-IR1*>+ (n=42). Statistical analysis: Kruskal-Wallis test X^2^=15.38 \*\*\**p* < 0.001. Pairwise comparisons were performed using Dunn’s multiple comparisons test. Statistically significant differences are represented by *p<0.05, ***p<0.001. ***C***, *dsxGal4>UAS-Dicer2*, SPR-IR1 (n=23); *dsx*Gal4>+ (n= 23); UAS-*Dicer*2, *SPR-IR1*>+ (n= 22). \*\*\*\**p* < 0.0001 One-way ANOVA with Tukey’s *post hoc* tests. ***D***, Mated females: *ppk*-Gal4>UAS-*Dicer*2, *SPR-IR1* (n=72); *ppk*-Gal4>+ (n=71); UAS-*Dicer*2, *SPR-IR1*>+ (n=62). Statistical analysis: Kruskal-Wallis test X^2^=107.5 \*\*\*\**p* < 0.0001. Pairwise comparisons were performed using Dunn’s multiple comparisons test. Statistically significant differences are represented by *p<0.05, ****p<0.0001. ***E***, Virgin females: *ppk*-Gal4>UAS-*Dicer*2, *SPR-IR*1 (n=41); *ppk*-Gal4>+ (n= 23); UAS-*Dicer*2, *SPR-IR*1>+ (n=45). \**p* < 0.0210 One-way ANOVA with Tukey’s *post hoc* tests. Dots represent independent flies, the mean and SEM are shown. n*s: not significant (p>0.05)*.

### LNvs are postsynaptic targets of PPK+ neurons

Our observations suggest that SP signaling through SPR expressed mostly in PPK+ neurons alters the temporal organization of rest-activity cycles in mated females inducing the loss of morning anticipation, a feature regulated by the circadian clock. We next wondered how the signal provided by PPK+ neurons would reach the circadian network. In order to address this question we first analyzed the expression pattern of the *ppk*-Gal4 driver in adult female by expressing a membrane-bound green fluorescent protein (UAS-mCD8-GFP) under the control of *ppk*-Gal4. We observed mCD8-positive expression in the reproductive tract, in the abdominal ganglia as well as in a few somas in the dorsomedial region of the central brain (**Figure 3**). Then, to explore whether PPK+ neurons could provide direct input to circadian neurons, we employed the anterograde trans-synaptic tracing tool trans-Tango (28) to define specific downstream synaptic partners of PPK+ neurons. Expression of the tethered trans-Tango ligand in neurons trigger mtd-Tomato expression in postsynaptic targets. We coexpressed the trans-Tango ligand and GFP to label pre-synaptic PPK+ neurons, and mtd-Tomato in the whole brain, in males, virgin and mated females in the event there were sexual and/or postmating differences in postsynaptic connectivity. Trans-synaptic labeling revealed that PDF-positive ventral lateral neurons, both the l-LNvs and s-LNvs, are postsynaptic targets to PPK+ cells in the three experimental groups (**Figure 4 and SF2**). We also observed a clear postsynaptic mtdTomato labeling in several unidentified neurons within the suboesophageal ganglia in male and female brains (**Figure 4 and SF2**). The subesophageal ganglia is a region within the CNS that is likely to contain circuits that mediate behavioral responses to mating. In addition, it has been reported that PPK+ projections are in close proximity to the somas of DN1 clock neurons, which are required to control the timing (onset) of the siesta in response to temperature (29). Although, trans-tango provided evidence of direct synaptic contact between PPK+ and PDF neurons, it failed to detect postsynaptic labeling within DN1 somas as previously reported (29), despite additional –yet unidentified-postsynaptic targets became apparent at the dorsal protocerebrum.

**Figure 3.**
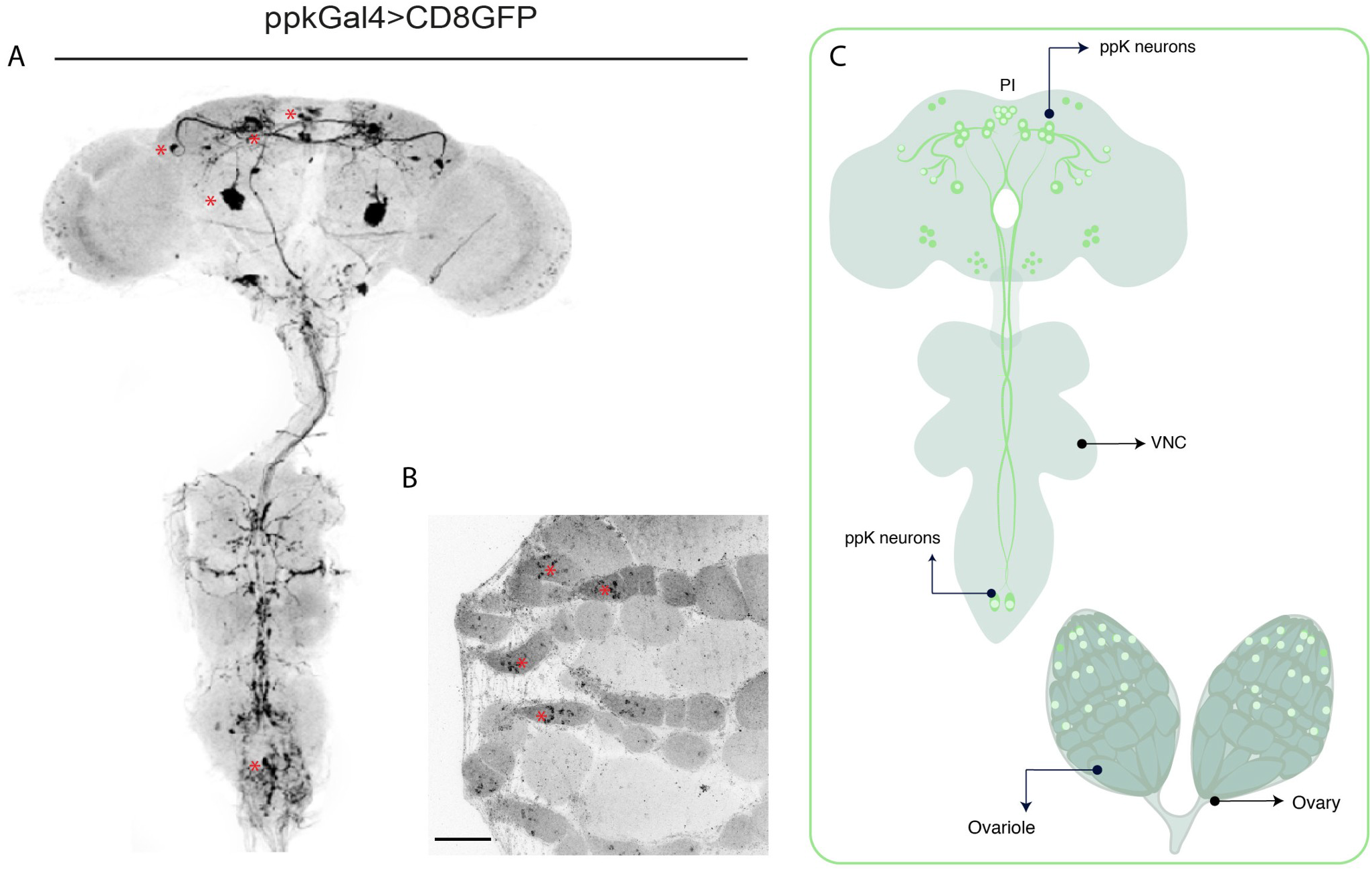
Expression pattern of the *ppk*-Gal4 line used in this work. (***A, B***) Confocal immunostaining of a representative brain and ventral nerve cord (***A***), and ovaries (***B***) of *ppk*-GAL4>UASmCD8GFP females stained with anti-GFP (green). Asterisks indicate the positions of stained PPK+ somas. Scale bar 100 μm for ovaries. (C) Schematic diagram showing the location of *ppk+* neurons.

**Figure 4.**
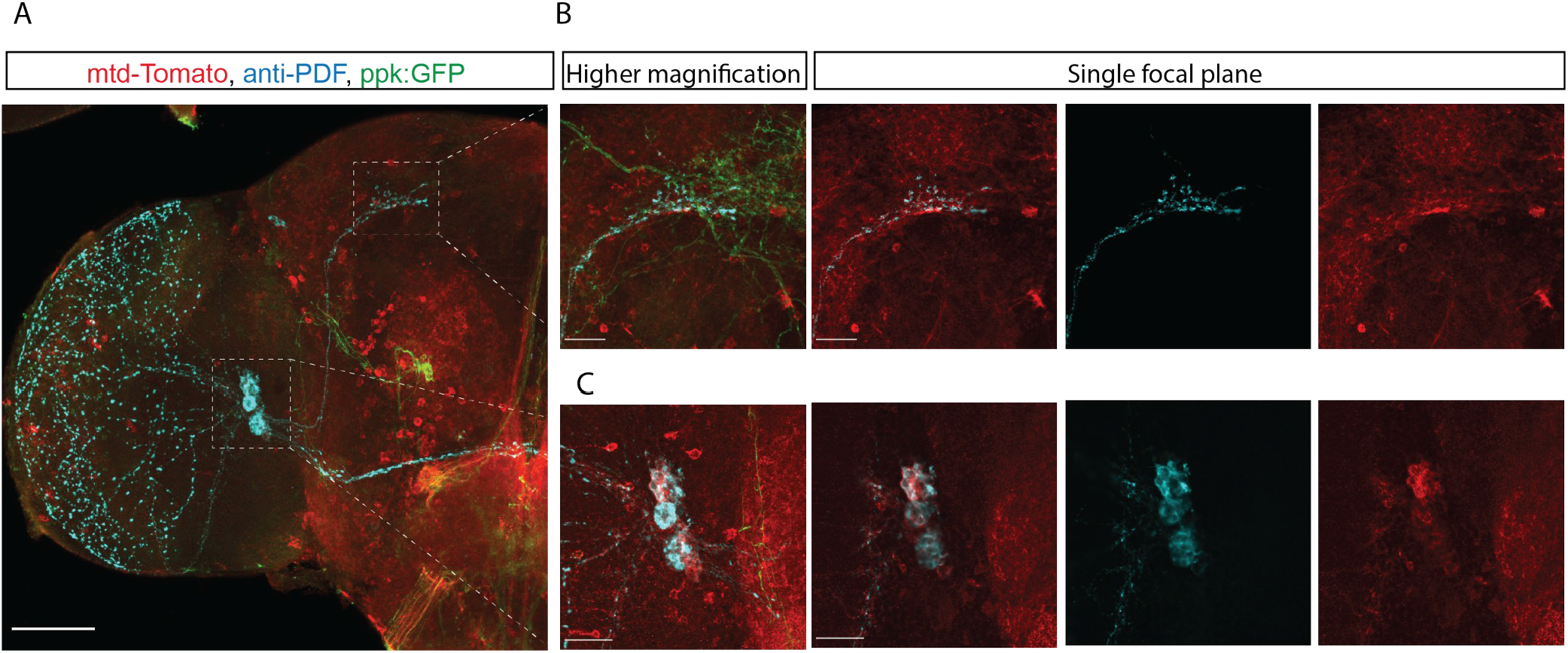
*pdf+* LNv neurons are postsynaptic targets of *ppk+* neurons. Trans-synaptic labeling using *trans*-Tango. *A*, Representative confocal image of a *ppk*-GAL4>UAS-myr-GFP, QUAS-mtdTom; trans-Tango mated female brain. Brains were co-stained with anti-PDF (cyan) to identify PDF+ neurons, anti GFP (green) to highlight *ppk+* presynaptic neurons and anti-DsRed (red) to identify putative *ppk* postsynaptic partners. The bar indicates 60 μm. (*B, C)* Higher magnification of the dorsal (*B*) and accessory medulla (*C*) regions of a mated female brain. The rightmost image of B and C panel shows single focal planes of an overlap between presynaptic and postsynaptic PPK contacts at PDF-expressing neurons in the accessory medulla (C) and in the dorsal termini of PDF axons (B). Bars indicate 20 μm.

Overall, these results confirm a contact between PPK+ and PDF+ neurons suggesting a (potentially) direct interaction between circuits involved in reproduction and the circadian clock. We next inquired what would be the impact of this communication onto the circadian clock.

### Mating reduces PDF levels

The sLNvs and ILNvs are required for coherent circadian behavior under free running conditions and are the only neurons within the circadian network that express the PDF neuropeptide (4–6). PDF levels at the dorsal sLNv terminals as well as in the somas oscillates in a circadian fashion; at dawn, levels are high while at dusk they are low (30). PDF released from the sLNvs appears to be responsible for the maintenance of free-running activity rhythms (31). Ablation of the LNvs and/or PDF results in the loss of morning anticipation, an advance in the evening activity peak under LD cycles, and a decrease in the free-running period (4,6). Thus, we reasoned that the phenotype associated to the loss of morning anticipation in mated females could possibly involve alterations in PDF levels, cycling, or both. To test this idea, we evaluated PDF levels by performing immunofluorescence in whole brains of mated and virgin females as well as in males. **Figure 5 A-F** shows that PDF immunoreactivity in males and virgin females displays a normal cycling pattern; however, PDF levels at the axonal termini are particularly reduced in mated females, especially in the morning, (albeit some residual cycling is still observed), suggesting that the mating state alters the circadian modulation of PDF levels (**Figure 5G**). To confirm whether reduced PDF levels in mated females could be mediated by SP signaling we knocked-down SPR in *ppk+* neurons. **Figure 5H** show that this restores PDF cycling in mated females and causes a significant increase of PDF levels in the morning when compared to mated controls. In fact, **figure 5H** also shows that PDF levels are similar to those observed in matched genetic controls as well as in *ppkG4*>SPR-IR1 males. Additionally, we noticed that virgin females devoided of SPR in *ppk+* neurons displayed a significant decrease in PDF levels when compared to control virgin females. Overall, these results indicate that PDF levels are responsive to postmating regulation in females.

**Figure 5.**
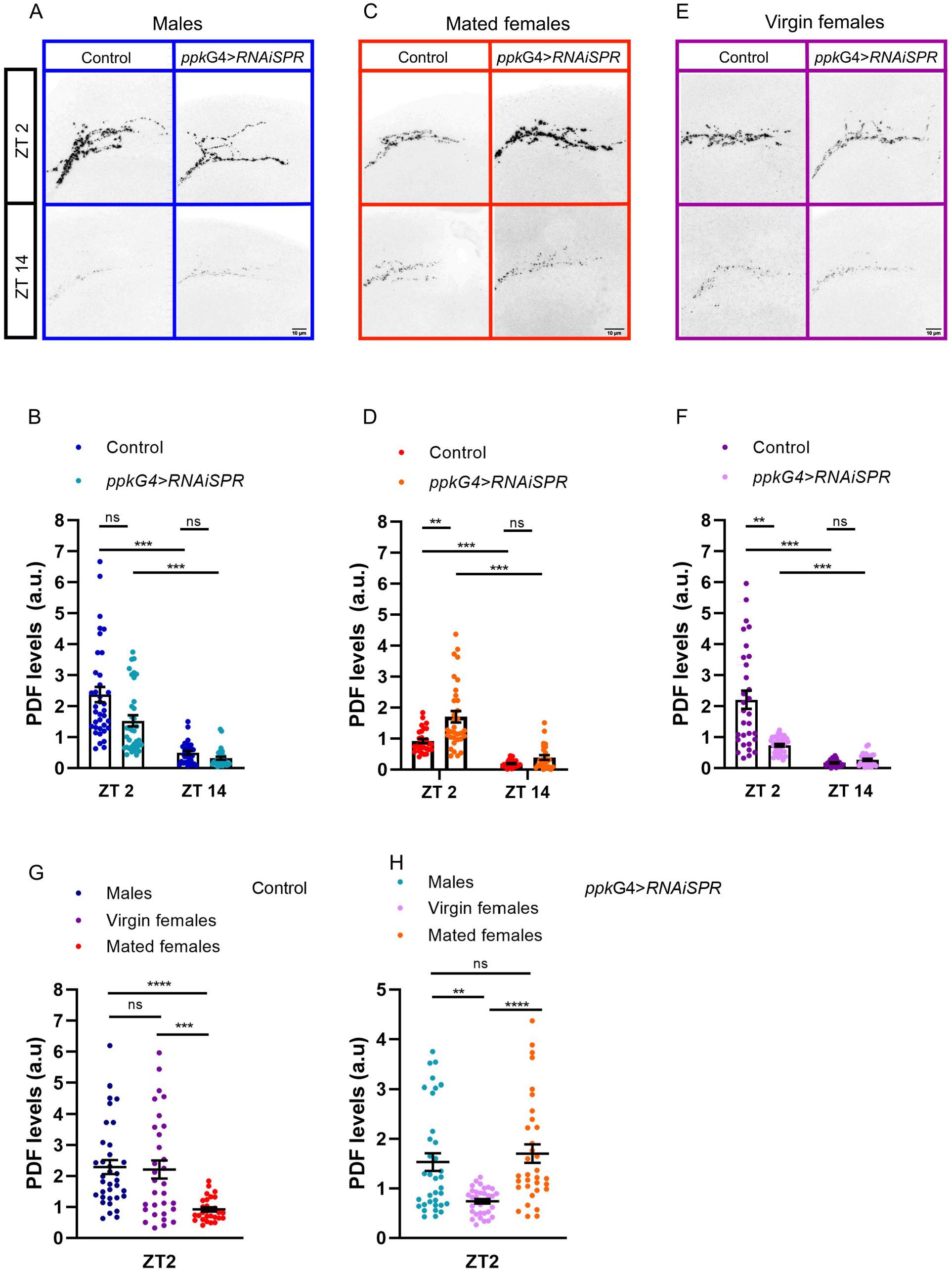
PDF levels are reduced in mated females. (***A, C, E***) Confocal images of representative sLNv dorsal projections of individual male, virgin and mated female flies showing their PDF content during the day (top) and night (bottom). Left: control flies (UAS-*Dicer*2, *SPR-IR1*>+). Right: flies with SPR downregulated in *ppk+* neurons (*ppk*-Gal4>UAS-*Dicer, SPR-IR1*). Brains were dissected at ZT02 and ZT14 and standard anti-PDF immunofluorescence detection was performed. The bar indicates 10 μm. **(*B, D, F)*** PDF quantitation of the sLNv dorsal projections for the twelve conditions mentioned above. Dots represent individual brains. ***B***, Males (n=31-37); ***D***, Mated females (n=26-33), ***F***, Virgin females (n=27-33). Symbols and error bars are average values +/− SEM. n*s: Not Significant (p>0.05)*, \*\**p* <0.01; \*\*\**p* <0.001; two-way ANOVA with Tukey’s *post hoc* tests. **(G and H)** Quantification of PDF signal intensity at day (ZT02) of control (UAS-*Dicer*2, *SPR-IR1*>+) and SPR downregulation in *ppk+* neurons (*ppk-*Gal4>UAS-*Dicer, SPR-IR1*) respectively. \*\*\*\**p* < 0.0001 Kruskal-Wallis test. In Dunn’s multiple comparisons test, statistically significant differences represented as **p< 0.01, ***p< 0.001, ***p < 0.0001. ns, not significant

## Discussion

In most animals, many behaviors display a high degree of temporal organization, even in the absence of any temporal cues from the environment, which is a clear indication of the presence of an endogenous circadian clock. Fruit flies have two daily bouts of consolidated activity, the first of which starts some time before lights on, under laboratory conditions, a property that is known as morning anticipation. However, after mating females display important changes in their behavioral repertoire, such as an increase in the oviposition rate and a reduction of daytime sleep (11,14). We show here that the temporal organization of locomotor behavior is also significantly altered after mating, since females loose their ability to prepare for the incoming morning (**Figure 1**). As it has been described with most postmating responses, such loss of anticipation can directly be related to the transfer of the sex peptide during copulation. It has been shown that many neurons have sex peptide receptors, which gives them the ability of initiating various postmating responses. Here we describe that the SPR+ *ppk*+ neurons are responsible for the suppression of morning anticipation. Furthermore, we show that the SPR activation in these neurons correlates with a decrease in the concentration of PDF, the main circadian neuropeptide, which is known to be actively involved in clock control of the morning peak of activity in *Drosophila*.

### Suppression of morning anticipation as a postmating response

During copulation, males transfer more than 100 proteins to receptive females (32), however SP is largely responsible for most postmating changes described in *Drosophila*. Most SP-dependent responses, including the loss of morning anticipation described here, are mediated by a small group of sensory neurons (SPSN) located within the female genital tract that co-express *ppk, dsx* and *fru* genes (17–19). However, the diverse properties of neurons that separately express *ppk, fru* or *dsx* make it unlikely that the SP response is only regulated by neurons that co-express the three genes. In fact, egg laying and receptivity are differentially affected by manipulations of subsets of these neurons (17–20,27,33). This appears to be the case with the loss of morning anticipation after mating. Indeed, our results show that this PMR is mediated mainly by the action of SP on *ppk+* neurons, since silencing SPR expression in them restores to a greater extent the ability to anticipate light/dark transitions in mated females. In SPSN, *fru*+ or *dsx*+ neurons, SPR downregulation achieved a much smaller effect, pointing to a more relevant SPR function in *ppk*+ neurons for this behavior (Figure 2). These results reinforce the idea that distinct SP target neurons regulate different postmating responses.

An intriguing question is how the signals triggered by mating reach higher-order neurons in the brain to coordinate the sensory integration of the postmating response. Mating signals can act by two major pathways (20,27). One is through SPSN neurons that can detect SP and alter its activation rate accordingly. This propagates the mating signal to the abdominal ganglia where, through direct synaptic contact, they silence the SAG neurons, which in turn relay mating information to the brain (17–19,27); alternatively, the mating signal could enter the hemolymph and act in higher centers of the brain (20). In this work we propose a new pathway through which SP could induce a postmating response; SP is detected by PPK+ neurons, which provide synaptic inputs to PDF neurons that drive clock-dependent morning anticipation (34). Interestingly, we observed this neuronal connection in female as well as in male brains, underscoring that this connection is not a sexually dimorphic feature (Figure 4 and Figure S2). The exact identity of the *ppk*+ neurons that are presynaptic to PDF neurons could not be determined since the *ppk*-Gal4 driver employed is expressed in the reproductive tract, in the abdominal ganglia as well as in some brain somas (Figure 3). Further work is necessary to pinpoint which specific *ppk*+ neurons contact PDF neurons directly.

### SP receptor plays a dual role in the temporal organization of locomotor behavior

The optimal time of day to engage in a particular behavior can vary depending upon environmental factors, and according to the internal physiology of the animal. In *Drosophila*, PDF differentially coordinates the activity of circadian neuronal groups to optimize behavioral output (5,7,34–36). PDF immunoreactivity at the axonal terminals of the sLNv oscillates in a circadian fashion (30). We show here that mated females have reduced levels of PDF compared with virgin females and males, and these levels can be restored when SPR expression is reduced in *ppk*+ neurons (Figure 5). This is expected because SP activates the inhibitory G-protein coupled receptor (GPCR) SPR in the SPSNs, which upon activation induces PMR by silencing the SPSNs directly and the SAG neurons indirectly (27). Consistent with this model, we hypothesize that PPK+ feed excitatory synaptic inputs to PDF neurons and the presence of SP silences PPK+ neurons, which in turn weakens or suppresses excitatory inputs to PDF neurons, inducing a decrease in PDF levels in the dorsal terminals (37). Further work will be necessary to address this hypothesis.

In virgin females PDF cycling/levels resembles the cycling shown in males; however, SPR downregulation in *ppk*+ neurons results in decreased PDF levels, in contrast to the effect observed in mated females (Figure 5). Although understanding this regulation is beyond the scope of the present manuscript, it is tempting to speculate that reduced SPR levels impairs the binding of additional neuropeptide/s that might also contribute to the neuronal communication among *ppk* and PDF neurons. This putative neuropeptide would likely be a GPCR ligand, just as SP is, but it may recruit different signaling pathways downstream of SPR. One attractive candidate is the myoinhibitory peptide (Mip), a potent agonist for SPR (38,39). Like other neuromodulators, Mip is highly pleiotropic, and it has been shown to modulate behaviours as diverse as sleep, feeding and mating receptivity (40–42). SPR mediates the sleep function of Mip but is dispensable for its feeding function and normal mating in virgin females (41,42). These findings should be thoroughly characterized in the future, as they suggest a different regulation of the excitability of sLNv through *ppk*+ neurons in virgin and mated females.

To the best of our knowledge, the loss of morning anticipation is the first postmating response that can be clearly ascribed to the circadian clock. The obvious question that arises is what could be the benefit of losing the ability to anticipate the end of the night. Along these lines, Fujii and collegues (2007) showed that male-female couples are highly active throughout the night and early morning, and this locomotor activity rhythm is associated with courtship (43). Thus, it is possible that, once mated, females lose the ability to anticipate dawn in order to reduce the chance of encounters with other males thus saving their energy for the daytime hours, which are probably better suited for feeding and oviposition activities, including searching for egg-laying sites. More generally, it is tempting to hypothesize that both the loss of morning anticipation and the suppression of the siesta could be part of a larger strategy of downregulating the circadian clock after mating, in order to restrict activity to daylight hours-whenever they come. Further confirmation of this hypothesis would require experiments to ascertain the preference for daytime over nighttime for the typical activities displayed by mated females (such as oviposition, feeding, searching for suitable egg-laying sites, etc.). To summarize, we show here that mating impairs the ability to anticipate dawn in females. This is probably achieved by SP-mediated silencing of *ppk*+ neurons, which in turn acts on the circadian release of PDF, thus obliterating a major signal that times locomotor activity to specific windows along the day. In nature, signaling of a wide range of sensory modalities along with internal cues must concomitantly be taken into account to match the onset of activity with environmental conditions. Our work provides a framework to unravel how mating triggered signals impinge upon clock neurons in the *Drosophila* nervous system and modifies the dynamic control of activity.

## Material and Methods

### Fly strains

All fly strains used in this study are detailed in **Table 1**. Flies were reared and maintained on standard cornmeal/agar medium at 25 °C and 60% humidity in a 12 hr:12 hr LD cycle unless stated otherwise. Six day-old adult males, virgin or mated females were used for all locomotor activity experiments.

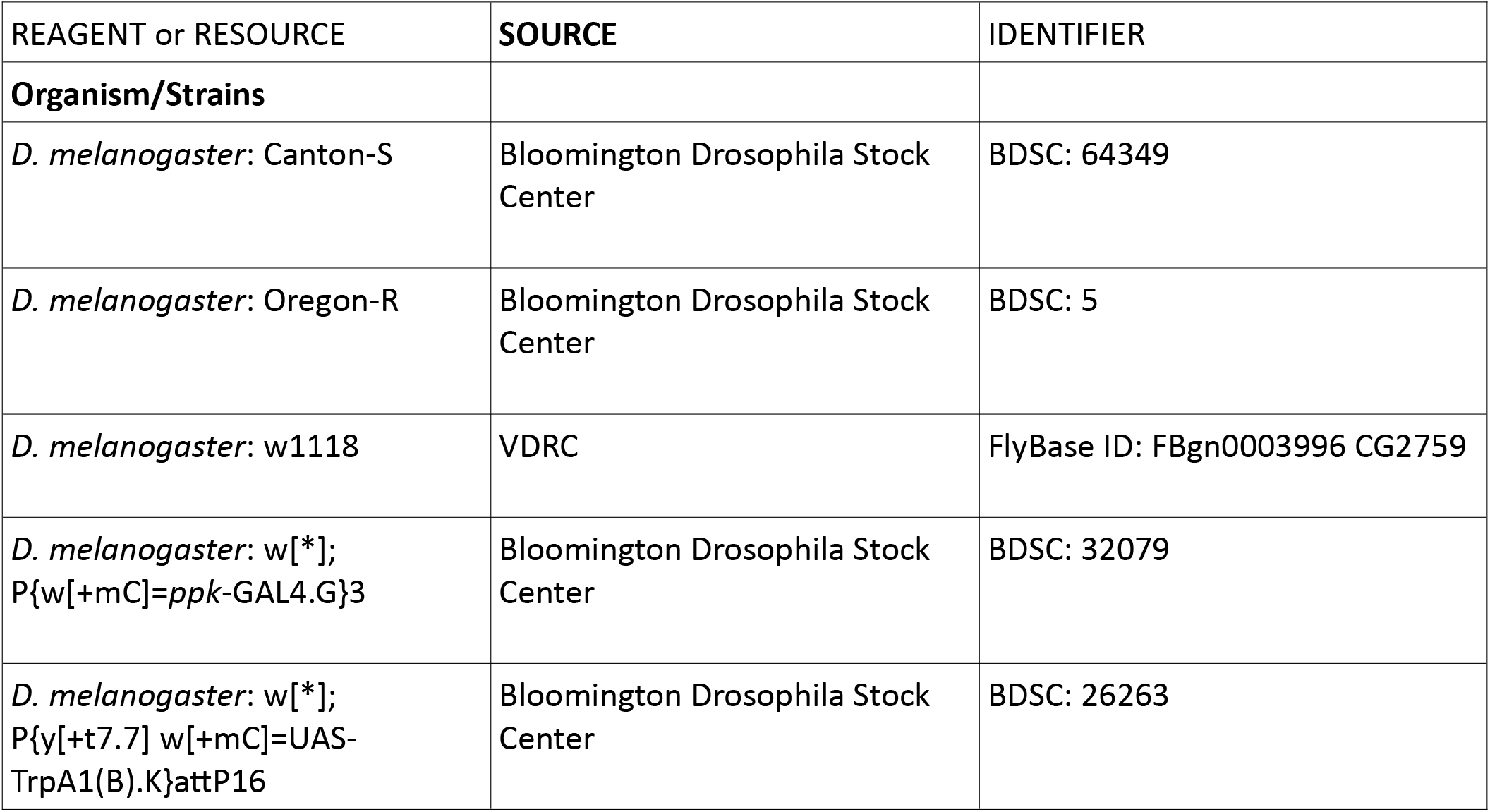
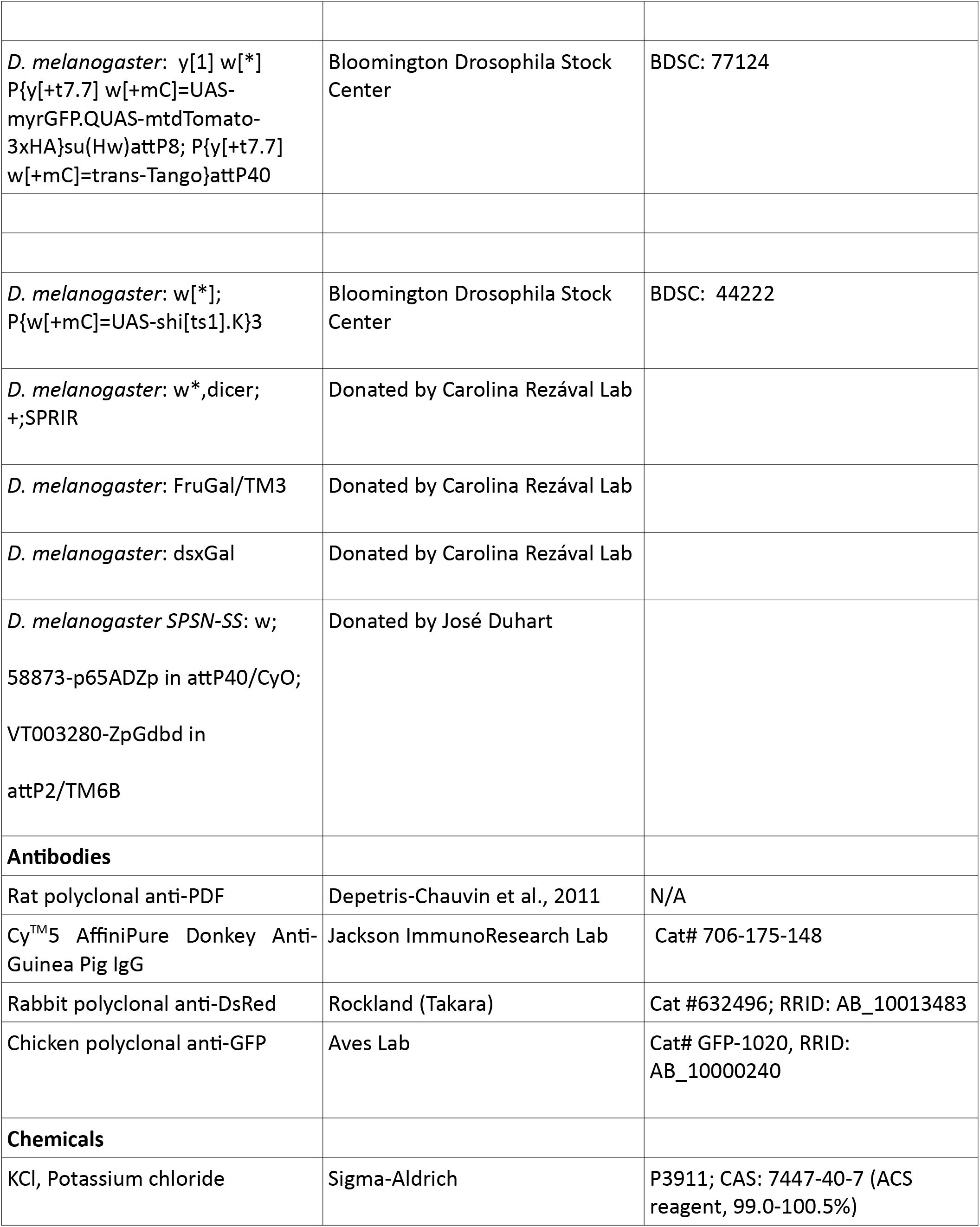
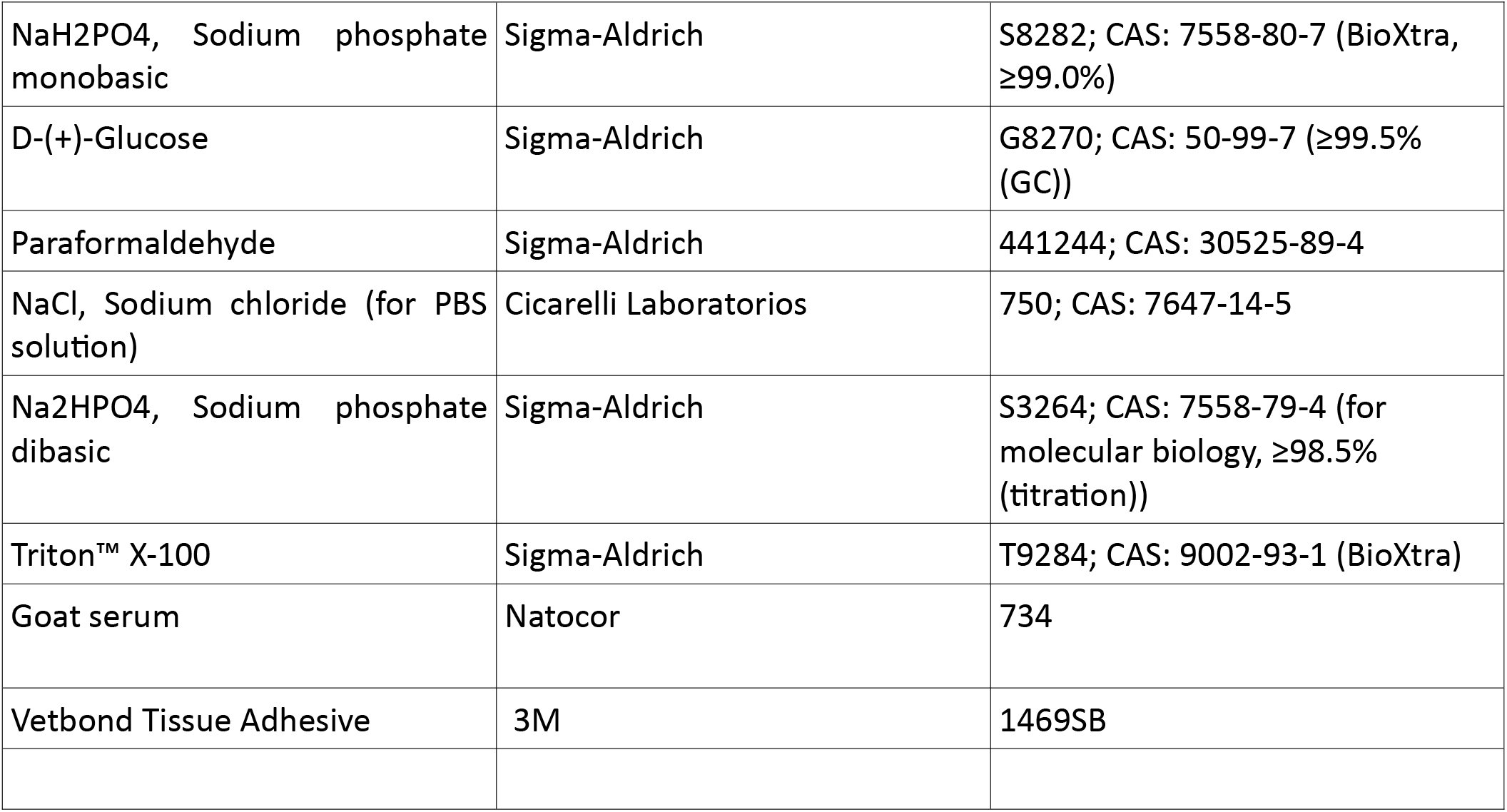

| REAGENT or RESOURCE | SOURCE | IDENTIFIER |
| --- | --- | --- |
| **Organism/Strains** |  |  |
| *D. melanogaster*: Canton-S | Bloomington Drosophila Stock Center | BDSC: 64349 |
| *D. melanogaster*: Oregon-R | Bloomington Drosophila Stock Center | BDSC: 5 |
| *D. melanogaster*: w<sup>1118</sup> | VDRC | FlyBase ID: FBgn0003996 CG2759 |
| *D. melanogaster*: w<sup>[*]</sup>; P{w[+mC]=ppk-GAL4.G}3 | Bloomington Drosophila Stock Center | BDSC: 32079 |
| *D. melanogaster*: w<sup>[*]</sup>; P{y[+t7.7] w[+mC]=UAS-TrpA1(B).K}attP16 | Bloomington Drosophila Stock Center | BDSC: 26263 |
| *D. melanogaster*: y[1] w[*]<br>P{y[+t7.7] w[+mC]=UAS-<br>myrGFP.QUAS-mtdTomato-3xHA}su(Hw)attP8; P{y[+t7.7]<br>w[+mC]=trans-Tango}attP40 | Bloomington Drosophila Stock<br>Center | BDSC: 77124 |
| *D. melanogaster*: w[*];<br>P{w[+mC]=UAS-shi[ts1].K}3 | Bloomington Drosophila Stock<br>Center | BDSC: 44222 |
| *D. melanogaster*: w*,dicer;<br>+;SPRIR | Donated by Carolina Rezával Lab |  |
| *D. melanogaster*: FruGal/TM3 | Donated by Carolina Rezával Lab |  |
| *D. melanogaster*: dsxGal | Donated by Carolina Rezával Lab |  |
| *D. melanogaster* SPSN-SS: w;<br>58873-p65ADZp in attP40/CyO;<br>VT003280-ZpGdbd in<br>attP2/TM6B | Donated by José Duhart |  |
| **Antibodies** |  |  |
| Rat polyclonal anti-PDF | Depetris-Chauvin et al., 2011 | N/A |
| Cy<sup>TM</sup>5 AffiniPure Donkey Anti-Guinea Pig IgG | Jackson ImmunoResearch Lab | Cat# 706-175-148 |
| Rabbit polyclonal anti-DsRed | Rockland (Takara) | Cat #632496; RRID: AB_10013483 |
| Chicken polyclonal anti-GFP | Aves Lab | Cat# GFP-1020, RRID:<br>AB_10000240 |
| **Chemicals** |  |  |
| KCl, Potassium chloride | Sigma-Aldrich | P3911; CAS: 7447-40-7 (ACS<br>reagent, 99.0-100.5%) |
| NaH<sub>2</sub>PO<sub>4</sub>, Sodium phosphate monobasic | Sigma-Aldrich | S8282; CAS: 7558-80-7 (BioXtra, ≥99.0%) |
| D-(+)-Glucose | Sigma-Aldrich | G8270; CAS: 50-99-7 (≥99.5% (GC)) |
| Paraformaldehyde | Sigma-Aldrich | 441244; CAS: 30525-89-4 |
| NaCl, Sodium chloride (for PBS solution) | Cicarelli Laboratorios | 750; CAS: 7647-14-5 |
| Na<sub>2</sub>HPO<sub>4</sub>, Sodium phosphate dibasic | Sigma-Aldrich | S3264; CAS: 7558-79-4 (for molecular biology, ≥98.5% (titration)) |
| Triton™ X-100 | Sigma-Aldrich | T9284; CAS: 9002-93-1 (BioXtra) |
| Goat serum | Natocor | 734 |
| Vetbond Tissue Adhesive | 3M | 1469SB |

### Video-based acquisition systems and locomotor behavioral analysis

Flies were entrained to 12:12h LD cycles during their entire development, and males, virgin and mated females were individually housed in open ended transparent chambers measuring 80 x 8 x 8 mm, containing approx. 1ml of banana medium (44) in one end and a cotton plug in the other end. A collective enclosure containing 26 of such chambers was built with two transparent acrylic boards (forming the floor and roof of all chambers) and white-opaque acrylic separations between chambers (to avoid visual contact between flies). The enclosures were placed inside an incubator (set at 25°C) over a translucid (not transparent) white plaque, and were illuminated from below with an array of white and infrared LEDs (850nm), in order to record activity both in LD and in constant darkness. Each enclosure was recorded from a distance of 18 cm, with two webcams whose infrared filters were manually removed.

Locomotor activity was monitored for 3-4 days in LD conditions. Data from the webcams was analyzed in a laptop computer (running Ubuntu, a Linux distribution) using custom made software (written in Python3 and using OpenCV libraries). The software implements a tracking paradigm: from the video signal the position of each fly is extracted in real time. The output is a text file containing a pair of coordinates (x, y) for each fly, sampled at one second intervals. These files are then processed with an analysis software we developed (in Bash) which provides statistics for activity (distance traveled, morning anticipation indexes, etc.) and sleep (daytime and nighttime sleep duration, number and duration of sleep bouts, etc.).

Sleep is defined as the absence of “significant activity” (defined as a displacement of at least one body length per second) for at least 5 consecutive minutes. The Morning Anticipation Index (MAI) is defined as the ratio between the total activity in ZT21-24 and the total activity in ZT 18-24 (26). Given there is a large variability in the MAI across different days in individual flies, we averaged the MAI of the first three complete days for every experiment for a more robust quantitation. All the software used is open-source and freely available upon request.

### Immunofluorescence detection

Adult flies were fixed with 4% p-formaldehyde (pH 7.5) for 60 min at room temperature. Brains were dissected and rinsed six times in PT buffer (PBS with 0.1% Triton X-100) for 30 min. Samples were incubated with rat anti-PDF (1:500; (45)) in 7% normal goat serum at 4°C for two days. Next, samples were washed in PT 6×10 min, and incubated with Cy5-conjugated anti-rat (1:500; Jackson ImmunoResearch, USA) for 2h at room temperature. Samples were washed 4×15 min in PT and mounted in Vectashield antifade mounting medium (Vector Laboratories, USA). For Trans-TANGO staining *ppk*-gal4 males were crossed with trans-TANGO females and kept at 25°C. Immediately after eclosion, adult males, virgin and mated females were separated from the progeny and aged for 15 days at 18°C. The immunofluorescence procedure was the same as described above, except for the length of the incubation with primary antibody, which was extended to 5 days at 4°C. The following primary antibodies were used: rabbit anti-DsRed (1:1000, Rockland), chicken anti-GFP (1:1000, Aves Labs) and rat anti-PDF (1:1000; (45)). The following secondary antibodies were used: Cy2-conjugated anti-chicken, Cy5-conjugated anti-rat, and Cy3-conjugated anti-rabbit (1:500, Jackson ImmunoResearch Laboratories, Inc). Images were acquired with a ZEISS LSM 880 Confocal Laser Scanning Microscope.

### PDF levels Quantification

For the quantification of PDF intensity at the sLNv projections, we assembled a maximum intensity z-stack that contains the whole projection (approximate 10 images) and constructed a threshold image to create a ROI to measure immunoreactivity intensity using ImageJ (NIH) (37). Data was analyzed with GraphPad Prism.

### Statistical analysis

The following statistical analyses were used in this study: one-way ANOVA and two-way ANOVA with post hoc Tukey’s or Holm-Sidak’s test for multiple comparisons of parametric data, and nonparametric Kruskal-Wallis statistical analysis with multiple comparisons as specified in figure legends. Parametric tests were used when data were normally distributed and showed homogeneity of variance, tested by D’Agostino & Pearson test. Sidak’s and Dunn’s multiple comparisons tests were performed after parametric and non-parametric ANOVA when GraphPad software was used. A p value < 0.05 was considered statistically significant. Throughout the manuscript n represents the total number of measurements compared in each experimental group (behaviour of an individual or brain), and N represents the number of independent times an experiment was repeated. In dot plots for MAI, daytime sleep and immunofluorescence quantification lines represent the mean value; error bars depict the standard error of the mean. No statistical methods were used to determine sample size. Sample sizes are similar to those generally used in this field of research. Samples were not randomized and analyzers were not blind to the experimental conditions.

In all the figures we show results of two or three independent experiments.

## List of non-standard abbreviations

DAM: Drosophila Activity Monitors
LD: Light-Dark conditions (12h light/12h dark)
MAI: Morning Anticipation Index
LNvs: lateral ventral neurons
lLNvs: large lateral ventral neurons
sLNvs: small lateral ventral neurons
PDF: Pigment Dispersing Factor
RNAi: RNA interference

## Acknowledgements

This work was supported by the Agencia Nacional de Promoción Científica y Tecnológica (Grants PICT-2016-1042 to S.R.G; PICT 2018-0995 to MFC and PICT-2019-091 to María Soledad Espósito and D.L.F.), the Universidad Nacional del Comahue, CRUB (Grant 04/B239 to D.L.F.). D.L.F., S.R.G and M.F.C. are members of the Argentine Research Council for Science and Technology (CONICET). SR and JJI hold graduate fellowships from CONICET. Stocks obtained from the Bloomington Drosophila Stock Center (NIH P40OD018537), were used in this study. We thank Carolina Rezával and Lucas Mongiat for helpful discussions and critical reading of the manuscript. We are also grateful to Carolina Rezával and José Duhart for sharing fly stocks with us.

## Supporting Information Captions

**Figure S1.**
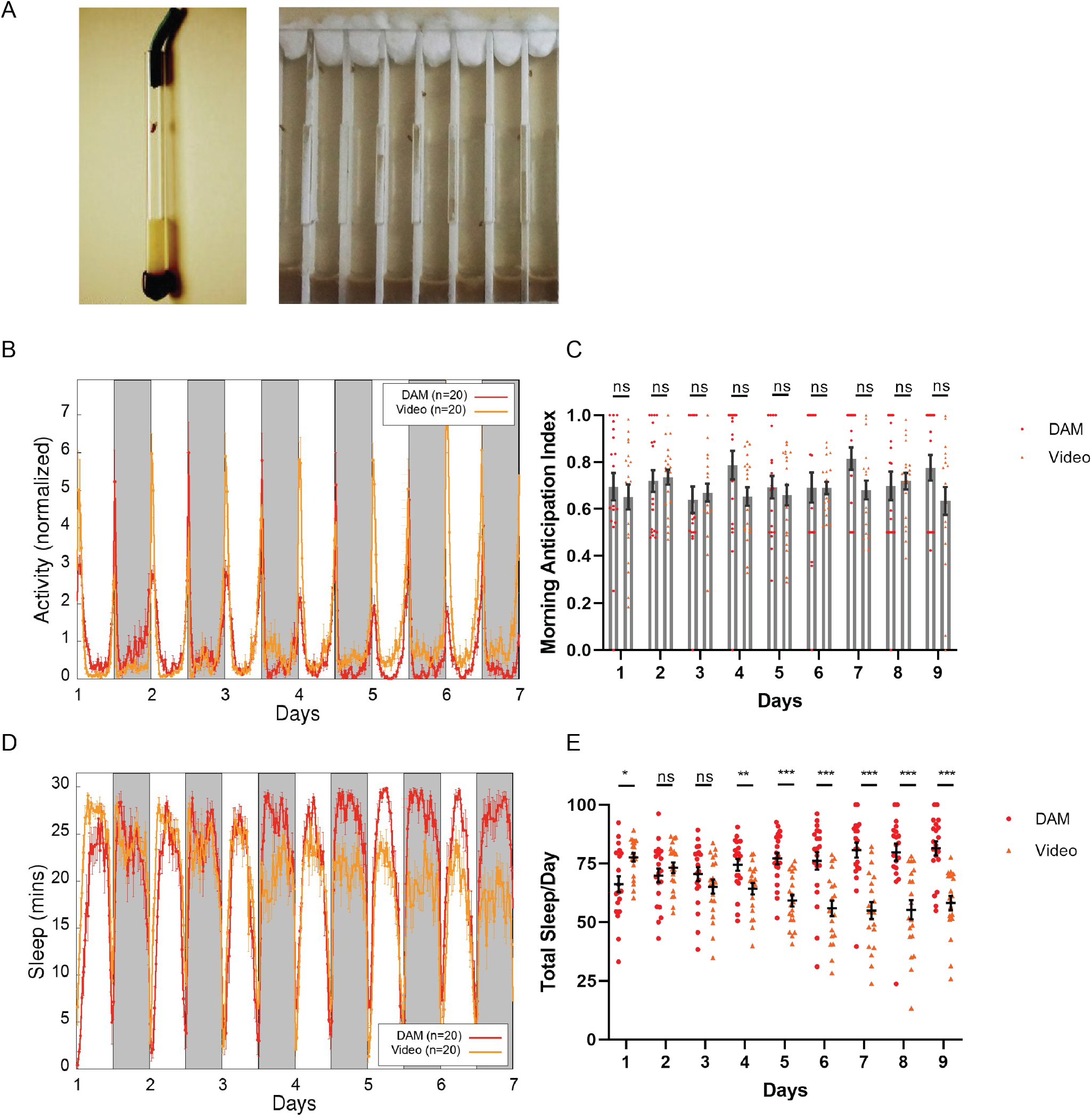
Comparisons between DAM and video acquisition systems. ***A*,** Left: Tube where a fly is housed in the DAM system. Right: Set of chambers where flies are housed in our video tracking system. ***B***, Comparison between group average locomotor activity of male *CantonS* flies in LD, obtained using both systems. ***C***, Comparison between Morning Anticipation Index of male *CantonS* flies, obtained using DAM (red) and Video sytems (orange). Each dot corresponds to the index calculated for a single fly. Statistical analysis: Scheirer–Ray–Hare test. Post hoc tests Wilcoxon rank tests for every day, corrected for multiple comparisons (Benjamini-Hochberg). ***D***, Comparison between the times spent sleeping (in 30 minutes bins) of male *CantonS* flies obtained using both systems under LD conditions. ***E*,** Comparison between total sleep of male *CantonS* flies obtained using DAM (red) and video (orange) systems under LD conditions. Each dot corresponds to the percentage of sleep per day calculated for a single fly. Statistical analysis: Scheirer–Ray–Hare test. Post hoc tests we have applied Wilcoxon rank tests for every day, corrected for multiple comparisons (Benjamini-Hochberg). Dots represent independent flies, the mean and SEM are shown Benjamini-Hochberg statistically significant differences *p< 0.05, **p< 0.01 ***p < 0.001. ns, not significant.

**Figure S2.**
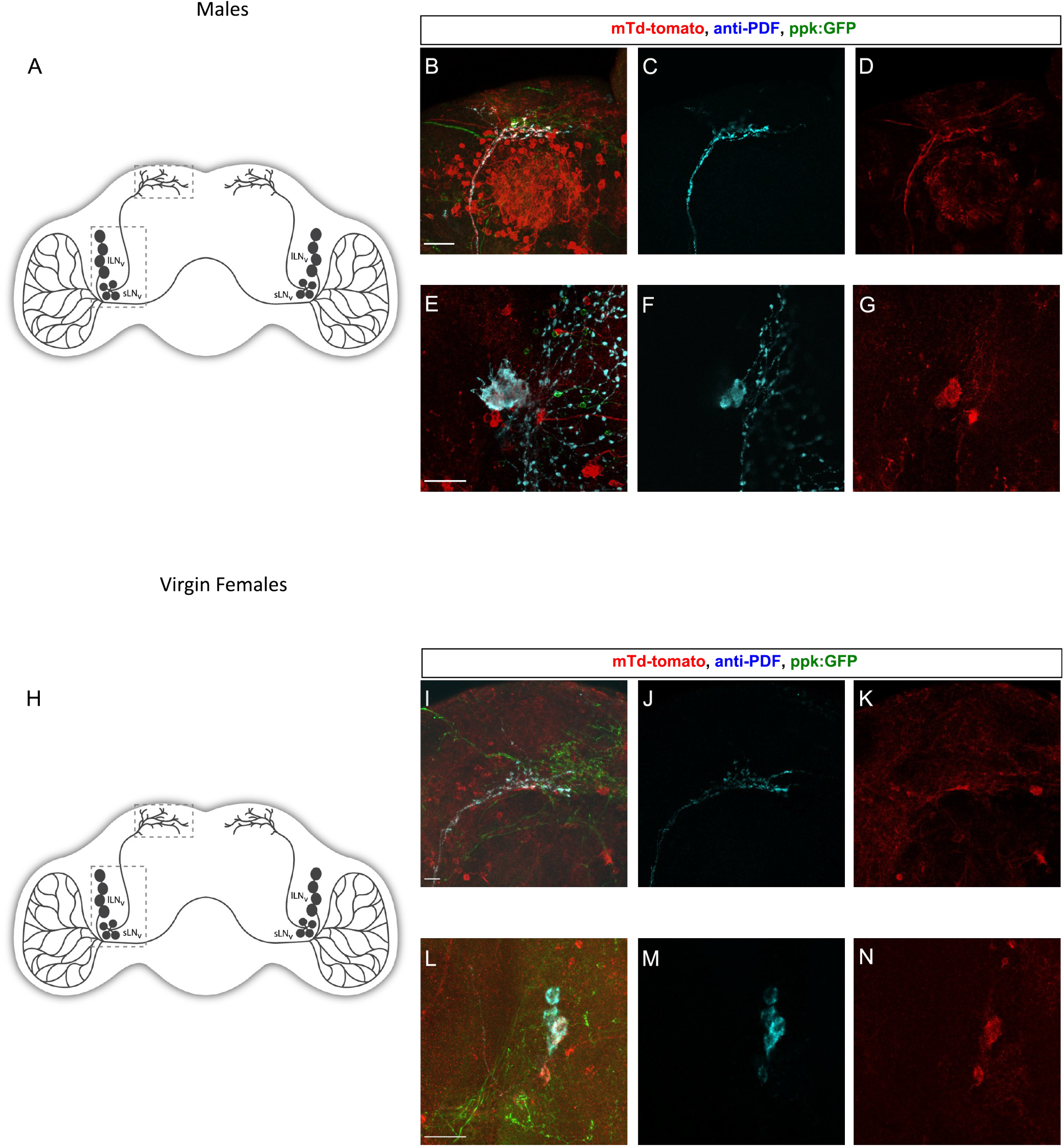
*pdf*+ LNv neurons are postsynaptic targets of *ppk*+ neurons in males as in virgin females. Trans-synaptic labeling using *trans*-Tango. ***A***, Schematic diagram of a *Drosophila* brain. ***B*,** Higher magnification of the dorsal region of *ppk*-Gal4>UAS-myr-GFP, QUAS-mtdTom; trans-Tango male brain. ***(C, D)*** Single focal plane of the image shown in ***B*** displaying the overlap of the PDF staining (cyan) with the postsynaptic *ppk*+ partners (red). ***E*,** Higher magnification of the accessory medulla region of *ppk-* GAL4>UAS-myr-GFP, QUAS-mtdTom; trans-Tango male brain. ***(F, G)*** Single focal plane of the image shown in ***E*** to exhibit the overlap of the PDF stain with the postsynaptic *ppk* partners (red). ***H*,** Schematic diagram of a fly brain. ***I*,** Higher magnification of the dorsal region of *ppk*-Gal4>UAS-myr-GFP, QUAS-mtdTom; trans-Tango virgin female brain. ***(J, K)*** Single focal plane of the image shown in ***I*** displaying the overlap of the PDF stain (cyan) with the postsynaptic *ppk* partners (red). ***L*,** Higher magnification of the accessory medulla region of *ppk*-Gal4>UAS-myr-GFP, QUAS-mtdTom; trans-Tango virgin brain. ***(M, N)*** Single focal plane of the image shown in ***L*** to show the overlap of the PDF stain with the postsynaptic *ppk* partners (red).

